# An integrated cardiac microtissue proteome map extends therapeutic remodelling by nanovesicles

**DOI:** 10.64898/2026.05.03.722552

**Authors:** Jonathan Lozano, Jarmon G Lees, Jonathon Cross, Anne M Kong, Ren J. Phang, Alin Rai, Shiang Y Lim, David W Greening

**Affiliations:** Baker Heart and Diabetes Institute, Melbourne, Victoria 3004, Australia; Baker Department of Cardiovascular Research, Translation and Implementation La Trobe University, Melbourne, Victoria 3086, Australia; O’Brien Institute Department, St Vincent’s Institute of Medical Research, Victoria, 3065, Australia; Department of Surgery and Medicine, University of Melbourne, Victoria, 3065, Australia; Drug Discovery Biology, Monash Institute of Pharmaceutical Sciences, Monash University, Victoria 3052, Australia; Baker Department of Cardiometabolic Health, University of Melbourne, Melbourne, Victoria, 3010, Australia; National Heart Research Institute Singapore, National Heart Centre, Singapore 169609, Singapore

**Keywords:** Cardiac microtissue, human pluripotent stem cells, endothelial, cardiomyocyte, fibroblast, nanovesicles, proteome remodelling

## Abstract

Human cardiac microtissues are a promising model to study cardiac biology and disease, but their application is constrained by therapeutic remodelling strategies and limited knowledge of their functional protein expression profiles. Here, we define the use of human cardiac microtissue (hCMT) model generated by assembling iPSC-derived endothelial cells, cardiac fibroblasts, and cardiomyocytes to model ischemia-reperfusion injury (IRI) through a model of hypoxia and reoxygenation and nanovesicle-mediated functional remodelling. Engineered nanovesicles (NVs), generated directly from human stem cells, have been shown to influence cardiac tissue and cell repair, and provide a platform for scalable and reproducible cell free-mediated therapy. We show the functional regulation of the hCMT model and define that administration of NVs (from human induced pluripotent stem cell origin) during reoxygenation significantly increase cardiomyocyte survival and preserve contractility function (contractile duration, relaxation time, relaxation:contraction velocity). Quantitative proteomics was applied to decipher the cell proteome dynamics and molecular mechanisms of IRI in our *in vitro* model following NV treatment, linked with networks associated with cell survival, energy production, and stress response regulation. Conserved proteome dynamics in NVs from different iPSC source reveal conserved upregulation of cellular protein networks involved in tissue repair (HSP70, CYFIP1), cardiac function (XIRP1, SLMAP, MYH6, CTNNA1, NDUFS2, GPD2), response to stress (CANX, PDCD6,), pro-survival (MDH2, LRPPRC, NIPSNAP1) and pro-angiogenic (FARSA, ECE1, RRAS) relative to vehicle treatments in context of IRI. Finally, we show that NVs also mediate differential remodelling in hCMT in response to IRI based on their cell origin, including altered wound healing and tissue repair response. Our findings provide an advanced human stem cell-based platform to understand underlying mechanisms of IRI and assess cell-free therapeutic cardioprotective strategies.

**Summary:** Advanced human stem cell-based platform provides a cardiac microtissue model to understand nanovesicle-based function and proteome remodelling, with potential applications for disease modelling and therapeutic intervention.

## Introduction

The human heart is an extraordinary organ composed of a diverse array of cell types, including abundant cardiomyocytes, fibroblasts, and endothelial cells, that function collaboratively to maintain its rhythmic contractile activity and systemic blood circulation^1^. This orchestrated interplay is critical for adapting to both physiological demands and pathological challenges^2^. However, when the coronary arteries are occluded, as in the case of myocardial infarction (MI), the resulting ischemia leads to cardiomyocyte death, triggering a cascade of inflammatory responses and fibrotic remodelling that irreversibly compromise heart function^3^. Despite advances in rapid reperfusion therapies that have emerged, many patients still develop long-term complications, including adverse ventricular remodelling and heart failure, as the regeneration of lost myocardium remains extremely limited in the adult heart^4,5^. Importantly, reperfusion injury following ischemia (due to the sudden tissue reoxygenation, termed IRI) exacerbates the initial injury and induces further cell death^6–8^.

The difficulty of studying and modelling the human heart to evaluate and develop advanced therapeutics underscores the need for standardized, representative in vitro models of human cardiac biology. Given that multiple cell types are involved in MI-induced acute cardiac injury and together they build a signalling network which drives tissue remodelling, *in vitro* modelling warrants the mimicking of cellularity and integration to be physiologically representative^9^. This versatile and dynamic network cannot be recapitulated entirely in two-dimensional (2D) monolayer culture. In this regard, scaffold-free tissue engineering approaches offer unique opportunities for developing three-dimensional (3D) models of the heart muscle in a microtissue (MT) structure^10,11^. In this format, cardiomyocytes can be seeded alone or in combination with other cardiac cell types, allowing cell aggregation and subsequent tissue formation, and mimicking the native physiological state^12–15^. Here, we defined the use of human cardiac microtissue (hCMT) model from endothelial, fibroblast and cardiomyocyte cells from induced pluripotent stem cell origin. These multicellular cardiac microtissues spontaneously contract and exhibit key features of cardiomyocytes, cardiac fibroblasts, and endothelial cells, including organized sarcomeres, calcium handling, and electrical activity, an intrinsic complexity to model ischemic reperfusion injury (IRI)^12,16^.

To limit the acute injury response following MI damage, regenerative strategies have garnered significant pre-clinical and clinical interest, including cell-based and cell-free therapies^17^. Although a variety of cell-based approaches have shown potential in preclinical studies, challenges such as poor cell retention, immune rejection, and the risk of arrhythmogenesis have hindered their current clinical feasiblity^18,19^. As a cell-free therapeutic strategy, nanosized cell-derived extracellular vesicles (EVs) have emerged as promising cell-free therapeutic strategy for cardiac repair^20^ due to their biophysical features in cardiac protection, feasibility in modification to enhance their bioactivity and extend circulation time, and derived from different stem cell sources^21–23^. Further, these stem cell-derived EVs can induce potent reparative effects to tissues and cells^24–27^ ^28^, can be targeted for their delivery and retention to the heart^29^, and recently EVs from regenerating tissue have been shown to induce protective and regenerative mechanisms directly to the heart^20^. As EV-like mimetics, nanovesicles (NVs) are sourced directly from stem cells, and overcome challenges in using natural EVs in their scalable generation, yield, rapid and reproducible generation, customizable engineering for bioactivity and targeted delivery, while still retaining natural cell-like properties like low immunogenicity and relevant surface proteins for function, making them superior for scalable drug delivery^30^. We have previously reported stem cell-derived NVs for cell-mediated uptake and their functional capacity^31,32^ and to promote cardiomyocyte survival, endothelial cell angiogenesis and attenuate primary cell fibrotic response^33^.

We have previously described the generation of 3D cardiac aggregates (microtissue)^12^ using human iPSC-derived cardiac cell types^16,34^. Integration of multiple cardiac cell types in these 3D cultures has been shown to promote the expression of cardiomyocyte maturity markers and better recapitulate *in vivo* microenvironment^35^. This multi-cell cardiac system exhibits spontaneous contraction, measurable extracellular field potentials, dynamic calcium cycling, and respond appropriately to chronotropic agents such as isoprenaline and carbachol^12,16,34^ . These features make hCMTs a reliable platform for modelling complex cardiac pathologies, including IRI^16,36^, diabetes-associated metabolic stress^37^ ^34^, and doxorubicin-induced cardiotoxicity^12^ . Furthermore, the hCMTs have proven effective in evaluating therapeutic interventions, such as metformin for diabetes-associated metabolic stress^37^ and mesenchymal stromal cell-based therapies for IRI^36^, highlighting their translational relevance for disease modelling and drug testing. These characteristics highlight the advantages of cardiac microtissues over traditional 2D cultures. Here, we aimed to employ a hCMT model featuring multiple and abundant cell types of the human heart (endothelial, fibroblast and cardiomyocyte cells) with cardiac identity to model IRI using hypoxia and reoxygenation and assess NV-mediated functional and proteome remodelling. This work shows the potential of using NVs to trigger protective and regenerative mechanisms in the context of IRI on human cardiac microtissues.

## Results

### Modelling ischemia-reperfusion injury with human iPSC-derived cardiac microtissues

Cardiac cell differentiation was performed using the human iPS-Foreskin-2 (CL2) cell line^38^, generating cardiomyocytes, endothelial cells, and cardiac fibroblasts (via the epicardial lineage), as detailed in Experimental Procedures. We have shown that these cardiac microtissues display multicellular organization and beating frequencies characteristic of the developing human heart, with vascularization and interspersed interstitial fibroblasts/mesenchymal-like cells among the cardiomyocytes^12^. To construct the cardiac microtissues, enriched day-19 iPSC-derived cardiomyocytes (1.2×10^5^ cells/well) were seeded onto Matrigel-coated UpCell plates, and iPSC-derived endothelial cells (7×10^4^ cells/well) and iPSC-derived cardiac fibroblasts (1 ×10^4^ cells/well) then seeded 24 hr following.

These cardiac microtissue sheets were cultured (24 hours), transferred, embedded, and cardiac microtissues then embedded in growth factor reduced Matrigel and cultured in cardiac microtissue medium, designated as day 0 (media changed every 2-3 days), before 7-10 days prior to ischemic conditioning and any functional assessments performed. An *in vitro* model of hypoxia/reoxygenation was developed, involving a sequential hypoxia incubation with ischaemic media, followed by reoxygenation with complete culture media, to simulate tissue injury caused by sudden reoxygenation (reperfusion injury). Normoxic hCMT (normoxia) and IRI cardiac microtissue (hypoxia-reoxygenation) conditions were then established and immunofluorescence confocal microscopy, cardiac troponin I (cTnI) quantification, and functional assessment analyses performed on this model (**Figure 1A**).

**Figure 1.**
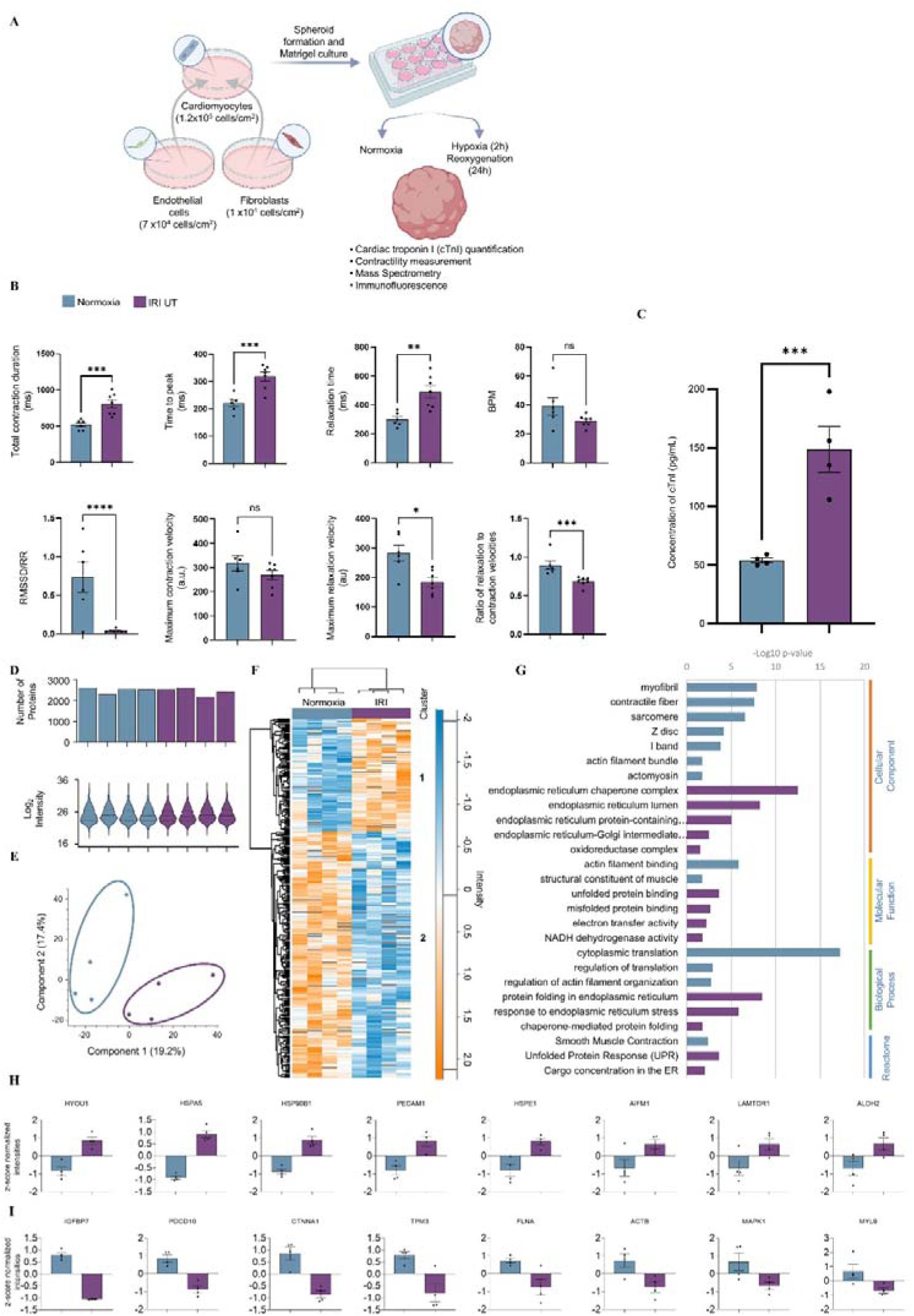
Human cardiac microtissue simulated IRI model. (**A**) Schematic overview of cardiac microtissue generation., culture, and hypoxia-reoxygenation workflow, with associated functional and analytical readouts. Human cardiac microtissue were generated from defined populations of human induced pluripotent stem cell-derived cardiomyocytes (60 %, 120,000 cells), endothelial cells (35 %, 70,000 cells), and cardiac fibroblasts (5 %, 10,000 cells). Cells were assembled as a detached cell sheet, transferred to suspension for spheroid formation, and embedded in Matrigel. IRI was modelled by simulated ischemia (2 hours, 1 % oxygen, N_2_-gassed ischaemic buffer) followed by reperfusion under normoxic conditions in complete microtissue media for 24 hours. Cardiac microtissues were assessed for cell viability, contractile function, and molecular changes using immunofluorescence and mass spectrometry-based proteomics. (**B**) Quantification of contractility parameters (normoxia n = 6, IRI n = 7). Error bars indicate S.E.M. (**C**) Cardiac troponin I (cTnI) release measured by ELISA (n = 4). Error bars indicate S.E.M. (**D**) Microtissue proteomic coverage and protein abundance (LFQ normalised intensity) for each group (n = 4). (**E**) Principal component analysis of cardiac microtissue proteome. (**F**) Hierarchical cluster analysis of normoxia and IRI microtissue samples identifies distinct protein expression profiles (ANOVA, p < 0.05, fold change ± 2.0). (**G**) Gene Ontology enrichment (cellular component, molecular function, biological process), and Reactome pathway enrichment analyses of differentially expressed proteins. (**H-I**) Representative proteins significantly upregulated (**H**) or downregulated (**I**) in response to IRI relative to normoxic/vehicle control. Based on -log10 p value. Error bars indicate S.E.M.

To visualise cell lineage markers associated in cardiac microtissues, immunofluorescence confocal microscopy revealed vimentin (a marker for fibroblasts), CD31 (a marker for endothelial cells) and cardiac troponin (a marker for cardiomyocytes) expression, which are the major cell types present in the myocardium *in vivo* (**Supp. Figure 1**). Contractile function assessment was then performed to assess functional impacts following IRI assay on cardiac microtissue. Our data highlight significant differences in total contraction duration (*p* < 0.001), time to peak (*p* < 0.0005), relaxation time (*p* < 0.005), maximum relaxation velocity (*p* < 0.05), ratio of relaxation to contraction velocities (*p* = 0.0005), and the Root Mean Square of Successive Differences (RMSSD) normalized to the R-R interval (i.e., RMSSD/RR) (*p* < 0.0001), commonly used in heart rate variability (HRV) analysis (**Figure 1B**). The latter being the clinical parameter which provides insights into the balance between the sympathetic and parasympathetic nervous systems, which can be affected by various factors like stress, exercise, and sleep^39^. Changes in HRV, including RMSSD, have been linked to cardiovascular health and other conditions^39,40^. To further evaluate the extent of tissue damage, cardiac troponin T (cTnT) release was performed, used clinically to detect suspected myocardial damage. hCMT subjected to IRI resulted in 3-fold increase in cTnT (*p* < 0.0005) in the conditioned media compared with normoxia controls (**Figure 1C**). These findings confirm that IRI of hCMT induces functional perturbation and cardiomyocyte death.

### Cardiac microtissue proteome remodelling with ischemia/reperfusion injury

To understand the cellular changes in human cardiac MT in response to hypoxia/reoxygenation, we performed proteomic profiling. Proteomic analysis was performed on cardiac MT in N = 4 during the culture period. Samples were lysed in SDS and reduced/alkylated before being subjected to protein hydrophobic and hydrophilic capture-based tryptic digestion^38^ and nLC-MS analysis. Proteins were subjected to direct data-dependent acquisition (DDA) MS combined with label-free quantitation^31,37^, processed through MaxQuant and analysed for reproducibility and quality assessment. In total, more than 2600 proteins were identified (**Supp. Table 1-3**). Of these identified proteins, 2396 proteins could be quantified across normoxia and hypoxic conditions; 2506 normoxia and 2505 in hypoxic condition (identified in at least two out of four replicates with maxLFQ algorithm) (**Figure 1D, Supp. Table 1-3**). Principal component analysis revealed the separation of cardiac microtissue proteomes in response to IRI modelling (**Figure 1E, Supp. Table 1-3**). To confer cardiac identity, we report 6/12 highly enriched heart tissue expressed proteins based on human protein atlas including actin and myosin binding proteins MYL4, MYH6, MYL7, MYBPC3, and heart muscle troponins TNNI3, TNNT2. We further report coverage of heart cell markers in this cardiac microtissue proteome, including cardiac fibroblasts (6 proteins; e.g., COL12A1, COL6A2, MRC2, FBLN1), cardiomyocytes (29 proteins; e.g., FABP3, MYL3, SDHA, TNNT2, MYBPC3), cardiac endothelial cells (1 protein; CD93), and cardiac smooth muscle cell protein MYH11 (**Supp. Table 1-2**).

We performed differential analysis of this cardiac microtissue proteome in response to IRI and detected 293 proteins that were significantly dysregulated in expression (normalized protein expression data, hierarchical K-means clustering, FDR_<_0.05, **Figure 1F, Supp. Table 4**; 92 upregulated, and 201 downregulated proteins. Importantly, we validate key cardiac cell markers in response to IRI indicating a remodelling and expression change induced by hypoxia in this assay, such as TPM1, MYL9, FLNA and MYOM1, cardiomyocytes markers (components of z-disc and contractile fiber), decreased in expression indicating a relative loss in cardiomyocyte composition/function. Gene Ontology-/Reactome-term annotation of the respective proteins and clusters revealed enrichment in normoxia proteome associated with cardiac structure (myofibril, contractile fiber, sarcomere, Z disc, I band, actin filament bundle and actomyosin), cardiac molecular function (structural constituent of muscle and actin filament binding), cardiac tissue regulation (regulation of actin filament organisation), and smooth muscle contraction (**Figure 1G**, blue bars, **Supp. Table 9**). In response to IRI, cardiac microtissue proteome cluster reflected enrichment in endoplasmic reticulum (ER) stress (ER chaperone complex, ER protein-containing complex, ER-Golgi intermediate compartment), oxidoreductase complex, networks associated with unfolded/misfolded proteins, electron/NADH transfer activity, and unfolded protein response (UPR) (**Figure 1G**, purple bars, **Supp. Table 9**). The data confirmed the patterns of an increase in intracellular stress networks, altered matrix protein networks and a decrease in cardiac cell composition linked with contractile functional protein networks in the heart with IRI.

Within these distinct cardiac microtissue proteome clusters in response to IRI, we identified several key proteins upregulated in expression associated with hypoxia response (HYOU1)^41^, heat shock proteins (HSPA5/90B1/E1)^42,43^, metabolic and energetic change (ALDH2, NDUFB3, HAGH, UQCRC2)^44–47^, and apoptosis (AIFM1)^48^ - previously attributed in response to myocardial IRI^33,40,49,50^ (**Figure 1H, Supp. Table 4**). In contrast, select proteins downregulated in response to IRI were involved in cell/tissue/matrix homeostasis, structural organisation, and contractile function (TPM1, CTNNA1FABP3, COL15A1, ACTB, VIM, COL18A1, MYL7/9/12A, MYOZ2, COLGALT1)^33,40,49,50^ (**Figure 1I, Supp. Table 4**). These findings demonstrate the patterns of an increase in hypoxia and stress response proteins, apoptotic regulators, and changes in metabolic and energetic protein networks, while a decrease in extracellular matrix organization, contractile response, and cytokine signalling proteins.

### Functional NVs protect human cardiac microtissues against simulated ischemia-reperfusion injury

To assess the therapeutic regulation of stem cell-derived nanovesicles in cardiac microtissue in response to IRI, we generated NVs from different iPSC lines and applied functional and proteome analyses to understand this system as a functional regulation platform. We have previously demonstrated that nanovesicles (NVs), generated directly from human iPSCs, influence cardiac tissue and functional repair mechanisms in 2D cardiac cellular models ^33^. NVs were generated from two different human iPSC lines (CERA and CL2), their membrane components was purified and biophysical and biochemical characterisation was performed based on their size, morphology and protein markers as previously described^33^ (**Figure 2A)**. Cryogenic transmission electron microscopy analysis revealed that particle size and distribution were similar between two cell sources (range 30-250 nm), with mean diameter of 86 nm (CERA) and 95 nm (CL2) (**Figure 2B-C**).

**Figure 2.**
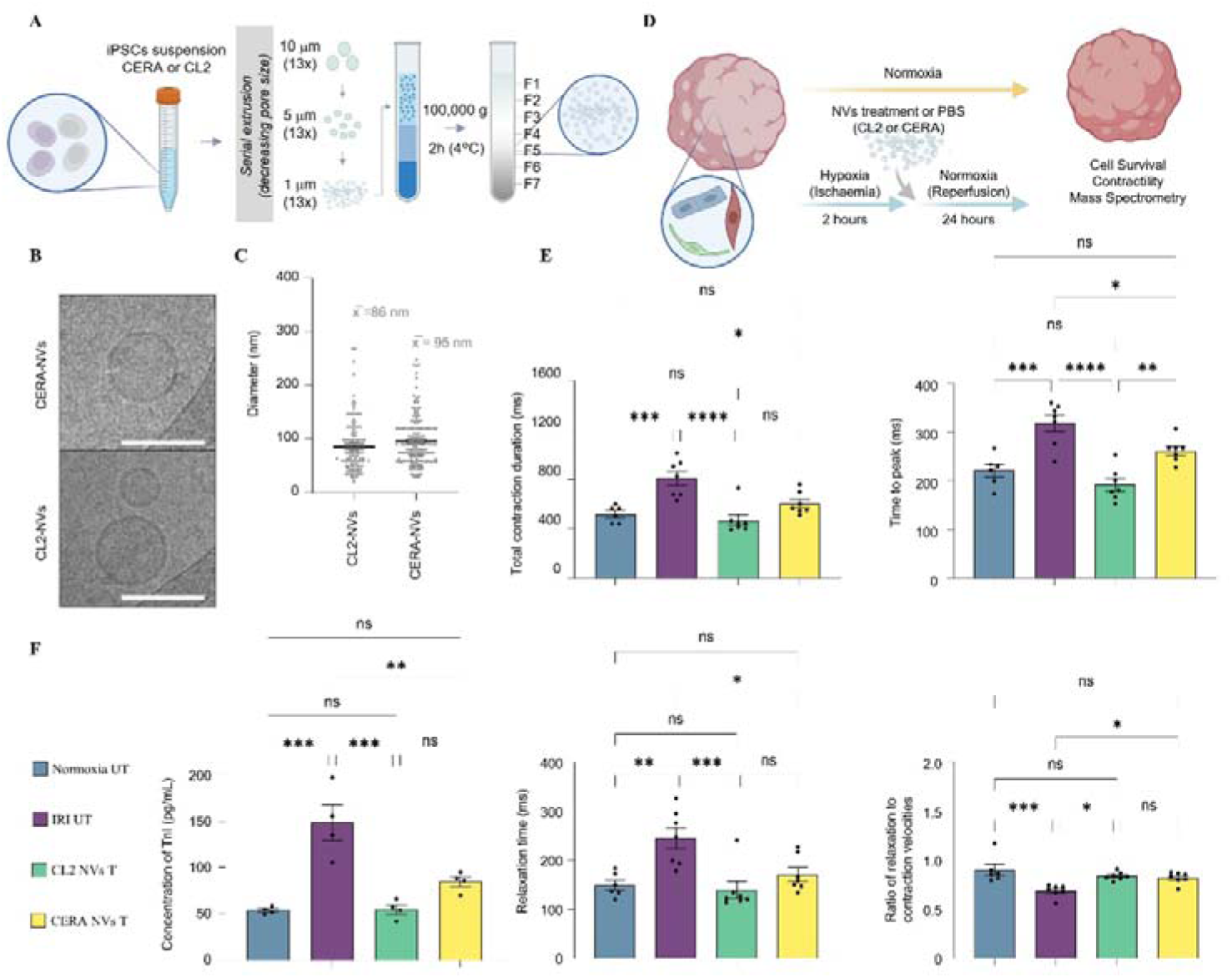
Generation of functional nanovesicles (NVs) from human iPSCs to protect against IRI-induced cell death and preserve contractility in human cardiac microtissues. (**A**) Human iPSC-derived NVs were generated by serial extrusion through 10, 5, and 1 µm polycarbonate membranes (13 extrusion cycles per membrane) and purified by density-cushion ultracentrifugation. The NV-enriched fraction (F5) was used for all subsequent biophysical characterisation and functional analyses. NVs were derived from two different hiPSC lines (CL2 and CERA). (**B**) Representative cryo-electron microscopy images showing spherical NV morphology. Scale bar = 100 nm. (**C**) Size distribution for NVs derived from CL2 and CERA iPSCs. (**D**) Experimental workflow for single-dose NV (or PBS) treatment of cardiac microtissues in the hypoxia-reoxygenation assay. Following 24 hours of reoxygenation, microtissues were analysed for cell viability, contractility and proteomic remodelling. (**E**) Quantification of contractility parameters in NV-treated groups (CL2 NVs T and CERA NVs T) and NV-untreated groups (Normoxia UT and IRI UT). Normoxia UT n = 6, all other groups n = 7. (**F**) Cardiac troponin I (cTnI) levels measured by ELISA (n = 4). Data presented as mean ± S.E.M (standard error of mean). \**p* < 0.05, \*\**p* < 0.005, \*\*\**p* < 0.001, \*\*\*\**p* < 0.0001, ns = non-significant.

Building on insights of proteome organization of our cardiac microtissue model in response to IRI, we aimed to define NV-mediated response in this model. Here, treatment groups included normoxia NV-untreated (normoxia UT, PBS/vehicle), and IRI groups: Hypoxia-reoxygenation untreated (hypoxic vehicle control: IRI UT), and NV treatment groups (CL2 NVs T and CERA NVs T). IRI groups were incubated for 2 hours in hypoxic media (N_2_ pre-gassed), followed by 24 hours in reoxygenation media supplemented with either NVs (single dose, 30 µg/mL, based on prior reported functional response in cardiac cells^33^) or vehicle control (PBS) **(Figure 2D)**.

To assess functional impact of NVs on hCMT model of IRI, contractility analyses was performed as previously described^12,16^. Here, we confirmed that our IRI model significantly impairs the functional capacity of hCMT compared to normoxia condition, including lengthening of total contraction duration (*p* < 0.001), extended time-to-peak (*p* < 0.0005), and increased relaxation time (*p* < 0.005). NV treatments mitigated these contractile perturbations induced by IRI (**Figure 2E**). Importantly, NV-treated hCMT exhibited relaxation-contraction velocity ratios comparable to normoxia controls, indicating improved diastolic (relaxation) function^51^ **(Figure 2E).** To evaluate whether NVs also provide pro-survival benefits, cTnT concentrations in conditioned media were measured. ELISA analysis showed that NVs from both CL2 and CERA iPSC lines significantly reduced cell death compared to vehicle-treated IRI group (CL2 NVs, *p* < 0.005, CERA NVs, *p* < 0.0005) (**Figure 2F**). These results confirm that distinct iPSC-derived NVs modulate the functional capacity and survival responses of hMCTs under hypoxic and reoxygenation conditions.

### NV-mediated remodelling of cardiac microtissue proteome in ischemia-reperfusion injury model

Cardiac microtissue proteomic response to NV treatment was performed. Here, after 24_hr following NV treatment in IRI assay (and vehicle hypoxic control), cardiac microtissue lysates were collected and proteome profiling performed. Proteome analyses revealed a network of proteins identified in each group and differential proteome clustering across groups confirmed by PCA and Pearson’s correlation analyses (**Figure 3A-D, Supp. Table 5**). Differential analysis by K-means clustering (*p* < 0.05) revealed 5 distinct clusters of protein networks based on expression in cardiac microtissue proteome, and a conserved NV differential response protein signature 58 in response to IRI (**Figure 3E, Supp. Table 5**).

**Figure 3.**
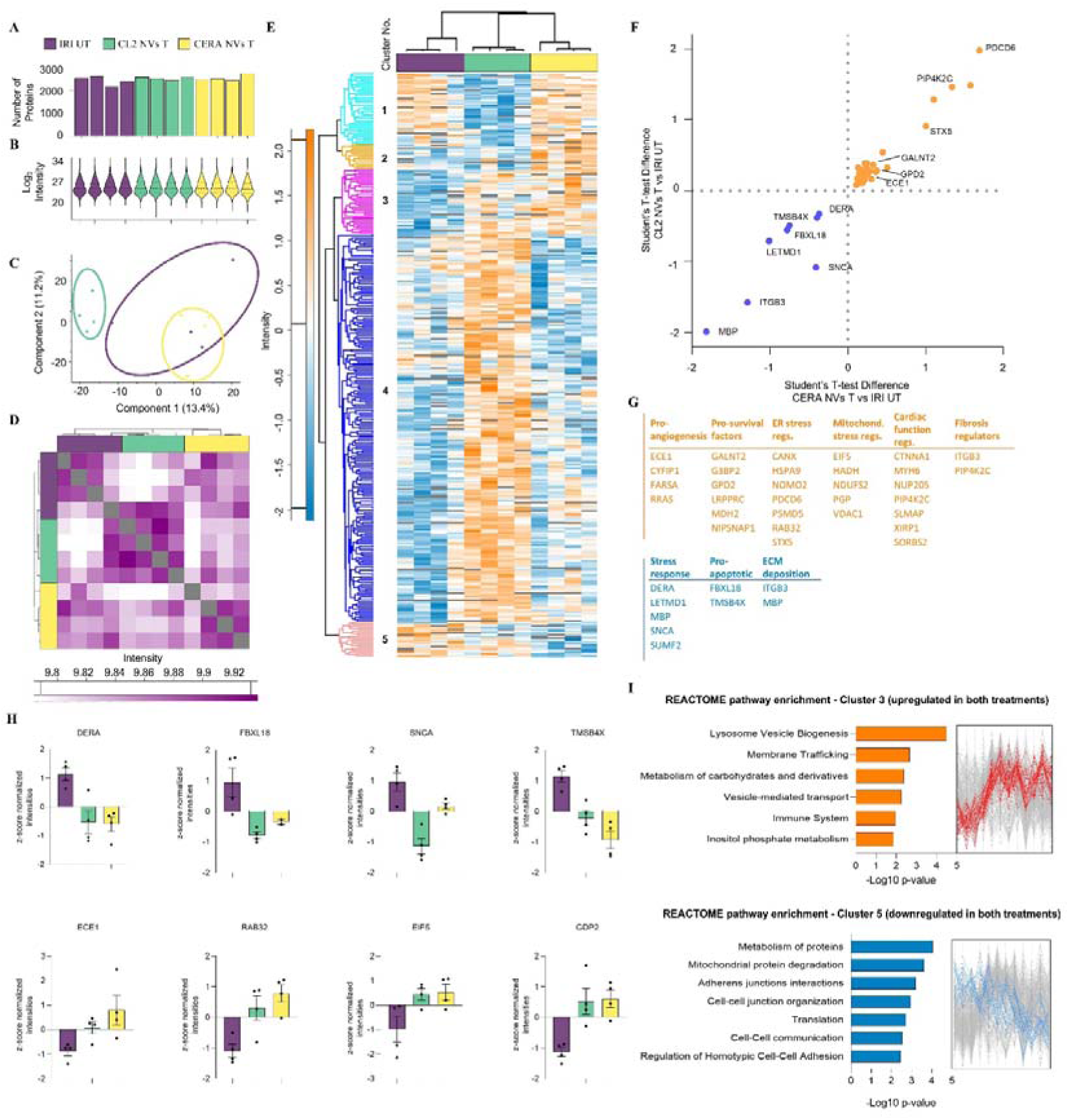
NVs were conserved in their proteome remodelling of cardiac microtissue model of simulated IRI model. (**A-B**) For organoid and NV-treated organoid groups, protein coverage/identification and abundance (LFQ normalised intensity) and (C) principal component analysis is shown (n = 4). (**D**) Pearson Correlation Analysis (R Score) of IRI UT, CL2 NVs T and CERA NVs T (n = 4). (**E**) Hierarchical cluster analysis of IRI UT, CL2 NVs T and CERA NVs T (*p* < 0.05, fold change ± 2.0), reveals 5 clusters of proteins differentially expressed. (**F**) Scatter plot for unique pro-reparative factors conserved in expression differences for both NV-treated groups. Student T-test difference, *p* < 0.05. (**G**) Differentially expressed proteins either up or downregulated following NV treatment that are associated with pro-reparative/wound healing function. (**H**) Differential protein abundance (fold change, of LFQ intensity, log2) of selected protein markers for each group, relative to IRI UT (control) (purple) (*p* < 0.05). Select proteins include pro-survival, pro-angiogenic, endoplasmic reticulum and mitochondrial stress regulators, and stress protein responders identified. (**I**) Enrichment analysis (ranked, *p* < 0.05) for Reactome pathway enrichment and protein intensity profile rank for Clusters 3 (upregulated in both treatments) and 5 (downregulated in both treatments).

We initially focused on NV-mediated remodelling conserved between both iPSC sources – i.e., CL2 and CERA NVs. Proteome analyses of cardiac microtissue proteome (NVs vs IRI) revealed statistically significant (p < 0.05) upregulation of proteins involved in angiogenesis (ECE1, FARSA)^52,53^, pro-survival (G3BP2, GPD2, RPPRC, MDH2, NIPSNAP1)^54–58^, endoplasmic reticulum stress regulation (HSPA9, NOMO2, PSMD5, RAB32, STX5)^59–63^, mitochondrial stress regulation (EIF5, HADH, NDUFS2, PGP, VDAC1)^64–67^, cardiac function (CTNNA1, MYH6, NUP205, PIP4K2C, SLMAP, SIRP1, SORBS2)^68–74^, and regulators of fibrosis (ITGB3)^75^ (**Figure 3F-H, Supp. Table 8 and 14**).

We observed further downregulated protein networks in response to NV treatments including key factors related to stress response (DERA, LETMD1, MBP, SNCA, SUMF2)^76–80^, apoptosis promoters (FBXL18, TMSB4X)^81,82^ and extracellular matrix deposition (MBP)^83^. GO-/Reactome-based enrichment analysis for cluster 3 (upregulated in both NV treatment groups relative to IRI vehicle) highlighted enrichment in networks associated with lysosome vesicle biogenesis, membrane trafficking, metabolism of carbohydrates and derivatives, vesicle-mediated transport, immune system and inositol phosphate metabolism (**Figure 3I**, top panel, **Supp. Table 10**). Downregulated protein cluster (cluster 5) showed enrichment in networks of cellular stress such as protein metabolism, mitochondrial protein degradation, regulation in adherens junctions, cell-cell junction organisation and cellular communication (**Figure 3I**, bottom panel, **Supp. Table 10**). Importantly, we showed that NVs mediate specific changes in cardiac marker expression in response to IRI, including upregulated expression of various cardiac associated signalling and structural networks proteins, including phosphatase PGP, ion channel VDAC1, translation factor DRG2 and dehydrogenase HADH. These results align with the functional data obtained further supporting the reprogramming capacity of functional NVs in cardiac microtissue.

### Selective NV reprogramming of cardiac microtissue proteome based on cell source

We next sought to elucidate differences in NV potency (**Figure 2E-F**) by examining proteome reprogramming induced by each NV source in the hCMT assay. Here, K-means clustering analysis revealed distinct cluster subsets for CL2-NV (**Figure 4A**) and CERA-NV (**Figure 4D**) relative to IRI vehicle control (**Supp. Tables 6 and 7**). Enrichment analysis revealed NVs from CL2 iPSCs (cluster 1) (**Figure 4B, Supp. Table 11**) involved in myofibril assembly, striated muscle cell development, cardiac (chamber/ventricle) muscle tissue morphogenesis, cardiac muscle contraction, and heart morphogenesis. Further enriched functions were associated with actin binding, structural constituent of muscle, unfolded protein binding and energy production (GTPase and ADP binding, pyruvate dehydrogenase activity, oxidoreductase activity). Reactome analysis showed aerobic respiration and respiratory electron transport, TCA cycle, regulation of apoptosis. Finally, we observed an enrichment in KEGG pathways involved in carbon metabolism, Citrate cycle (TCA cycle), and fatty acid degradation. Ontology analysis of cluster 2 (**Figure 4C, Supp. Table 11**) showed a downregulation of cellular components involved in focal adhesion and muscle thin filament tropomyosin, biological processes linked to actin filament organisation and mitotic spindle organisation, and molecular function in actin binding and pyruvate transmembrane transporter activity, with pathways analysis decreased in regulation of actin cytoskeleton.

**Figure 4.**
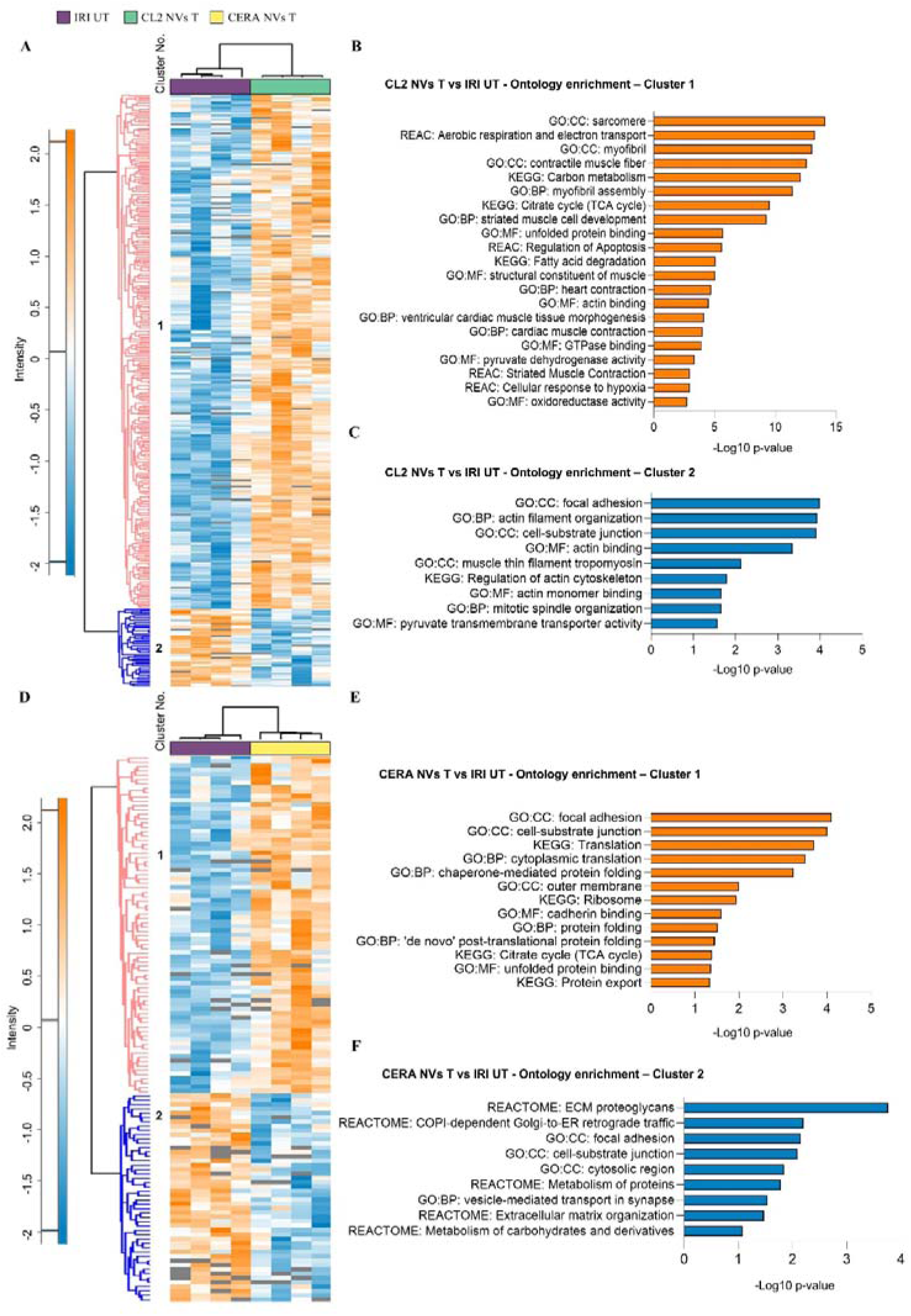
Human iPSC origin confers selective NV-mediated remodelling of cardiac microtissues. (**A**) Hierarchical cluster analysis of CL2 NVs T and IRI UT (*p* < 0.05, fold change ± 2.0), reveals distinct clusters of differential protein expression. (**B**) Functional enrichment analysis (ranked, *p* < 0.05) of CL2 NVs T and IRI UT for Reactome pathway enrichment, Cellular Component, KEGG pathways, Molecular Function, Biological Process for Cluster 1. (**C**) Enrichment analysis (ranked, *p* < 0.05) of CL2 NVs T and IRI UT for Cluster 2. (**D**) Hierarchical cluster analysis of CERA NVs T and IRI UT (*p* < 0.05, fold change ± 2.0). (**B**) Functional enrichment analysis s (ranked, *p* < 0.05) of CERA NVs T and IRI UT for Cluster 1. (**C**) Enrichment analysis (ranked, *p* < 0.05) of CERA NVs T and IRI UT for Cluster 2.

CERA iPSC-derived NVs (cluster 1) (**Figure 4E, Supp. Table 12)** revealed enrichment of pathways related to cytoplasmic translation and protein folding, including cadherin binding and unfolded protein binding. Intracellular networks were enriched for mitochondrial outer membrane, focal adhesion, and cell-substrate junction components, with functional associations linked to voltage-gated anion channel activity, ceramide binding, and metabolic response. In contrast, proteins downregulated in response to CERA-NVs treatment relative to the IRI vehicle control (cluster 2) (**Figure 4E, Supp. Table 12 and 13)** were enriched for focal adhesions, cell-substrate junctions, extracellular matrix organisation and proteoglycans, as well as metabolism of carbohydrates and derivatives.

These differences in NV-mediated proteomic remodelling of hCMT indicate that, while NVs from different iPSC lines broadly enhance functional capacity and promote a pro-reparative cardiac response, the extent and nature of this remodelling are cell-source dependent. Specifically, CL2-NVs reprogram hCMT in the IRI assay towards sustained basal cardiac function processes, corresponding with greater functional potency, whereas CERA-NVs predominantly modulate cellular stress responses and mitochondrial function networks.

## Discussion

Here, we describe a comprehensive analysis of proteins expressed by human iPSC-derived cardiac microtissue and tie these findings to proteome changes that occur relative to therapeutic intervention by NVs, fundamental knowledge needed to advance the use of cardiac microtissue to model human disease and molecular insights towards deliverable cell-free therapeutics. We demonstrate that NVs tune multiple pro-tissue repair networks, including cardiac structural and function, as well as regulators of cellular stress and metabolism^33,40,49,50^. While the therapeutic potential of these protein networks often relies on direct delivery to the heart to reactivate regenerative pathways in adult cardiomyocytes, NVs provide a potent and deliverable strategy to translate such regulation into effective, targeted therapeutics. Collectively, this work highlights the potential of human iPSC derived-NVs to activate protective and regenerative mechanisms in the context of IRI. Crucially, we extend our study beyond solely a description of the expression of these proteins by also demonstrating the functionality and relevance of protein networks in our microtissue model system to IRI in cardiac injury response.

Cardiac repair following MI remains a challenge in the clinic. Recent research has identified the need of a scalable multimodal approach given that the injury triggers simultaneously diverging signals known as cardiac remodelling. However, the study of the human heart represents a major hurdle to develop efficient therapies^84,85^. Engineered hCMT now represent mature, multicellular, human-relevant models that recapitulate structural integration, vascular-stromal crosstalk, and clinically meaningful functional responses under stress conditions such as IRI^86^. Recent work in this field spans tri-culture heart_on_a_chip platforms with iPSC_derived cardiomyocytes, fibroblasts, and endothelial cells that exhibit flow_responsive endothelium and catecholamine responses^86^; hCMTs that support electrical pacing, arrhythmia simulation, and mitophagy reporters for IRI studies^87^; and hCMTs that model acute MI and fibrotic remodelling under hypoxia-reoxygenation^3^. Collectively, these advances underscore that human microtissues can now reproduce hallmarks of IRI (contractile dysfunction, biomarker release, mitochondrial stress, and maladaptive extracellular matrix (ECM) responses) while enabling high-content, mechanistic analyses and therapeutic testing^3,86–90^.

Our study extends this trajectory by challenging iPSC_derived nanovesicles (NVs) in a human iPSC microtissue model of IRI^16,36^. The hCMT model used, as shown by Yin et al.^12^, recapitulates key structural and functional features of human cardiac tissue, including the integration of cardiomyocytes, fibroblasts, and endothelial cells and establishment of a communication network. We also demonstrate that in the absence of treatment, IRI induced marked contractile abnormalities such as prolonged contraction duration and time_to_peak, slowed relaxation, and a diminished relaxation/contraction velocity ratio that were accompanied by depressed beat_to_beat variability, a clinically used proxy of autonomic vagal/parasympathetic regulation in post_MI assessment^91^. These changes mirror clinical observations that heart rate variability declines after MI and correlates with recovery and risk, supporting the translational relevance of our readouts^91–95^. Biochemically, hCMT subjected to IRI microtissues exhibited a significant increase in cTnT release, consistent with the established role of cardiac troponins as biomarkers of cardiomyocyte injury and necrosis in MI^96,97^.

Our proteomics findings delineate a stress_response reprogramming under IRI. Control microtissues were enriched in proteins associated with cardiac structure and function, including sarcomeric components and actin-binding proteins, reflecting a homeostatic myocardial phenotype. In contrast, IRI microtissues exhibited upregulation of proteins involved in ER stress, unfolded protein response (UPR), and oxidative metabolism, including HYOU1^41^, HSPA5^42,43^, and AIFM1^48,98^, hallmarks of cellular stress and apoptosis. These findings are canonical molecular signatures of IRI pathobiology: ER stress sensors drive translational arrest, chaperone induction, and, when unresolved, apoptotic cascades^99^; the integrated stress response converges on eIF2α phosphorylation to modulate translation during ischemia^100^; and mitochondrial damage with impaired respiration and oxidative stress is central to reperfusion injury^101^. The concordance of our proteome and functional assessments with these mechanisms and previous reports^49,50,102^ validates the microtissue platform as a human surrogate for myocardial IRI.

Ideally for clinical utility under MI, a cardiac repair therapy will regulate simultaneously cell survival, angiogenesis, and scar formation. In this study, we show that following hypoxia a single dose of either iPSC NVs (CL2 or CERA) administered during reperfusion, increased the expression of pro-survival factors (G3BP2, GPD2, RPPRC, MDH2, NIPSNAP1)^54–58^ on the treated microtissues. This indicates preservation of cellularity count in the treated groups and further confirmed as a reduction of cardiomyocyte cell death measured as free-cTnI (*p=*0.005). These results extend previous findings in 2D cultures and underscore the therapeutic potential of iPSC-derived NVs in myocardial injury^33^. We show that NVs treatment ablate the disfunction induced by IRI in microtissue contractility by recovering basal levels of contraction duration, time to peak, relaxation time, and the ratio of relaxation to contraction velocities which aligns with the increase in the expression of proteins associated with cardiac function and contraction (CTNNA1, MYH6, NUP205, PIP4K2C, SLMAP, SIRP1, SORBS2)^68–74^ as revealed by mass spectrometry analysis. Furthermore, and in line with a multimodal approach for cardiac tissue repair, our proteomic analysis shows that NVs induce the expression of proteins associated with promotion of angiogenesis (ECE1, FARSA)^52,53^ while also modulating fibrosis regulators (ITGB3)^75^. Further functional analysis associated to these mechanisms is warranted. In addition, our proteomic reprograming analysis also shows that NVs recover the homeostatic levels of proteins associated with ER-stress (HSPA9, NOMO2, PSMD5, RAB32, STX5)^59–63^ and mitochondrial dysfunction (EIF5, HADH, NDUFS2, PGP, VDAC1)^64–67^. These processes have been shown to be hallmark to contend ischemia/reperfusion-induced damage. We propose that their regulation by NVs is key to preserve cellularity and function of the injured tissue. Further experiments to elucidate the mechanism and effect in vivo are warranted.

Interestingly, the proteomic signatures of NV-treated microtissues differed depending on the source of iPSC. Both CL2 and CERA are well-known somatic iPSC lines used in cardiac differentiation studies. CL2 iPSC^38^s were generated from human neonatal foreskin fibroblasts using viral transfection to mediate cellular reprogramming, whereas CERA iPSCs^103^ were produced via a specific non-integrating reprogramming method. Here, CL2 NVs were potent in their remodelling function, and promoted enrichment in pathways associated with cardiac development, myofibril assembly, and contractile function, suggesting a reprogramming toward restoration of baseline cardiac physiology. In contrast, CERA NVs favoured pathways related to mitochondrial integrity, cellular stress response, and membrane dynamics, indicating a more adaptive response to injury. This divergence and their background origin/origin reprogramming strategy, highlights their distinct functional utility towards tailoring NV-based therapies based on desired reparative outcome, and potential therapeutic use.

In summary, these results demonstrate that NVs mediate select, potent effect in CMT platform during reoxygenation that is sufficient to preserve contractility, promote cardiomyocyte survival, and reprogram their proteome toward cardiac repair and stress resilience. We highlight cell source impacts generation of NVs and should lead to further studies employing other stem cell or immune cell sources in their generation to fine tuning selective tissue repair response. Interestingly, these differences did not impact significantly the function of the microtissues at the evaluated timepoints. These findings position iPSC-derived NVs as promising candidates for regenerative therapies and cardiac microtissue platforms to assess and screen potential IRI-based therapeutics^104^.

## Experimental Procedures

### Cell culture

#### Human induced pluripotent stem cells (iPSCs)

Human iPS-Foreskin-2 (CL2) cell line^38^, kindly provided by James A. Thomson (University of Wisconsin), and CERA007c6 (CERA)^103^ iPSC line were maintained on vitronectin-coated plates in TeSR-E8 medium (Stem Cell Technologies, VA, USA) according to the manufacturer’s protocol. Briefly, cells were cultured until confluent, then detached using an enzyme-free dissociation reagent (ReLeSR, Stem Cell Technologies). The resulting cell aggregates were re-seeded onto new vitronectin-coated plates for further expansion. Medium was replaced every 2-3 days. Brightfield images were obtained using an inverted microscope (Olympus IX71, Tokyo, Japan) at 40x magnification.

#### Cardiomyocyte differentiation from iPSCs

Cardiomyocytes (CMs) were derived from human iPSCs as previously described with modifications^36,105,106^. Briefly, human iPSCs were seeded onto human embryonic stem cell qualified-Matrigel (Corning) coated plates at a density of 1.25×10^5^ cells/cm^2^ in TeSR-E8 medium supplemented with 10 μM Y−27632 (Abcam). After 48 hours (day 0), medium was replaced with RPMI-1640 basal medium (Thermo Fisher Scientific) containing B-27 without insulin supplement (Thermo Fisher Scientific), growth factor reduced Matrigel (1:60 dilution) and 10 μM (for CL2 iPSCs) or 6 μM (CERA iPSCs) CHIR99021 (Cayman Chemical). At day 1, medium was replaced with RPMI 1640 basal medium containing B-27 without insulin supplement. At day 2, medium was changed to RPMI 1640 basal medium containing B-27 without insulin supplement and 5 μM IWP-2 (Sigma-Aldrich) for 72 hours. From day 5, cells were cultured in RPMI 1640 basal medium containing B-27 supplement (Thermo Fisher Scientific) and 200 μg/mL L-ascorbic acid 2-phosphate sesquimagnesium salt hydrate (Sigma-Aldrich) (cardiomyocyte medium, v), with CMm changed every 2-3 days. At day 12, differentiated cardiomyocytes were dissociated into single cells and split 1:4 onto growth factor reduced Matrigel-coated plates in DMEM/F-12 GlutaMAX medium supplemented with 20% FBS (Sigma-Aldrich), 0.1 mM 2-mercaptoethanol, 0.1 mM nonessential amino acids, 50 U/mL penicillin/streptomycin and 10 μM Y−27632. At day 13, medium was changed to CMm. From days 14–19, CMs were enriched to >95% cardiac troponin T positive cells by culturing in glucose-free DMEM medium (Thermo Fisher Scientific) supplemented with 4 mM lactate (Sigma-Aldrich).

#### Endothelial cell differentiation from iPSCs

Human iPSCs were differentiated into CD31+ endothelial cells according to a previously published method^16,36,107^. For endothelial differentiation, iPSCs were dissociated into single cells and seeded onto growth factor reduced Matrigel-coated plates at a density of 1×10^5^ cells/cm^2^ in TeSR-E8 medium supplemented with 10 μM Y−27632. After 24 hours, referred to as day 0, medium was replaced with DMEM/F12 GlutaMAX medium (Thermo Fisher Scientific) containing N-2 supplement (Thermo Fisher Scientific), B27 supplement, 8 µM CHIR99021 and 25 ng/mL BMP4 (STEMCELL Technologies) for 3 days. Medium was then replaced with StemPro-34 SFM complete medium (Thermo Fisher Scientific) supplemented with 200 ng/mL VEGF-165 (Peprotech) and 2 µM forskolin (Sigma-Aldrich) for 3 days. At day 6, CD31 positive cells were sorted by FACS using a CD31-conjugated antibody (BD Pharmingen), and expanded on human fibronectin (Merck) coated plates and cultured in EGM2-MV medium (Lonza) supplemented with 50 ng/mL VEGF-165.

#### Cardiac fibroblast differentiation from iPSCs

Human iPSCs were differentiated into cardiac fibroblasts according to a previously published method^16,35,36^. iPSCs were dissociated into single cells and seeded onto Matrigel-coated plates at a density of 2.5×10^4^ cells/cm^2^ in TeSR-E8 medium supplemented with 10 μM Y−27632. After 24 hours, referred to as day 0, medium was replaced with STEMdiff™ APEL™ medium (STEMCELL Technologies) containing 20 ng/ml BMP4, 20 ng/ml Activin A (Peprotech), and 1.5 µM CHIR99021 for 3 days. On day 3, medium was replaced with STEMdiff™ APEL™ medium containing 30 ng/ml BMP4, 1 µM retinoic acid (Sigma-Aldrich), and 5 µM IWP-2 (Sigma-Aldrich), and then replaced again on day 6 but without IWP-2. On day 9, cells were dissociated and reseeded at 1.5×10^4^ cells/cm^2^ on human fibronectin in STEMdiff™ APEL™ medium containing 10 μM SB431542 (Tocris Bioscience) and 10 μM Y−27632. On day 13, epicardial cells were dissociated and reseeded at 2.5×10^4^ cells/cm^2^ on 0.1% porcine gelatine-coated plates and cultured in STEMdiff™ APEL™ medium containing 10 ng/ml bFGF (Peprotech) and 10 μM Y−27632 and medium was changed every 2 days without Y−27632. On day 20, the medium was changed to FGM-3 (Lonza) and replaced every 2-3 days until confluent.

#### Engineered cardiac microtissues

To construct the cardiac microtissues, as previously described^12,16,37^, enriched day-19 iPSC-derived cardiomyocytes (1.2×10^5^ cells/well) were seeded onto Matrigel-coated 48-well NuncTM UpCell plates in DMEM/F-12 GlutaMAX medium supplemented with 20% fetal bovine serum (Sigma-Aldrich), 0.1 mM 2-mercaptoethanol, 0.1 mM nonessential amino acids, 50 U/mL penicillin/streptomycin and 10 μM Y−27632. After 24 hours, iPSC-derived endothelial cells (7×10^4^ cells/well) and iPSC-derived cardiac fibroblasts (1 ×10^4^ cells/well) were then seeded onto the iPSC-cardiomyocytes and cultured in cardiac microtissue medium consisting of a mixture of CMm, EGM2-MV medium, and FGM-3 medium (at 1:1:1 ratio) supplemented with 50 ng/mL VEGF-165. After 24 hours, the UpCell plates were brought to room temperature, and the detached cell sheet was transferred to an ultralow attachment plate (Sigma-Aldrich) containing cardiac microtissue medium for 24 hours. The resulting spheroids were then embedded in 15 µL of growth factor reduced Matrigel and cultured in cardiac microtissue medium, designated as day 0. Cardiac microtissues were maintained in culture in a humidified CO2 incubator on an orbital shaker at 100 rpm and medium was changed every 2-3 days.

#### Immunofluorescent staining of engineered cardiac microtissue

Engineered cardiac microtissues were fixed in 10% neutral buffered formalin for 1 hour at room temperature and then dehydrated in 20% sucrose solution for 24 hours. Dehydrated samples were embedded in Optimal Cutting Temperature compound (Sakura Finetek, Tokyo, Japan) and cryosections (6 µm thick) were treated with 0.2% Triton X-100 permeabilization buffer and Protein Block (Dako, Victoria, Australia) before staining with primary antibodies; cardiac troponin T (2 μg/mL, rabbit polyclonal, Abcam) and CD31 (4 μg/mL, mouse monoclonal, clone JC70A, Dako) or vimentin (0.3 μg/mL, mouse monoclonal, clone V9, Dako), followed by Alexa Fluor-488-conjugated goat-anti-rabbit (10 μg/mL, Invitrogen) and Alexa Fluor-594-conjugated goat-anti-mouse (10 μg/mL, Invitrogen) secondary antibodies. Sections were then counterstained with 1 μg/mL of DAPI (Invitrogen) for nuclear staining. Epifluorescence images of immunostained sections were acquired with an Olympus BX61 upright microscope using analySIS software.

#### Human iPSC nanovesicle generation and purification

Generation of nanovesicles (NVs) was performed as described^108^ with modifications. Briefly, adherent iPSCs (CL2 and CERA) were individually harvested (following PBS wash) using a solution of EDTA (10 mM) (3×3 min round incubation) and cell suspension spun at 500 × g for 5 mins. The pellet was resuspended in PBS and the cell suspension sequentially extruded through 10, 5, and 1 µm polycarbonate membranes (19 mm; Advanti Polar lipids, 610010) (13 times across each filter, Whatman). The extruded NVs were subsequently purified using 10% OptiPrep™ (Stemcell Technologies) density cushion (step gradient formed by overlaying extruded sample on 10% and 50% iodixanol) and centrifuged at 100,000 × g for 2 hr at 4_°C. Seven equal fractions were obtained, diluted in PBS (to 1.5 mL), centrifuged at 100,000 × g for 2 hr at 4_°C (TLA-55 rotor; Optima MAX-MP ultracentrifuge) and resuspended in PBS and stored at -80 °C until further use. The yield (protein) and density of each fraction was determined as described^109^.

For comparison, natural extracellular vesicles (EVs) from each iPSC line were obtained as described previously^109^, based on isopycnic (iodixanol density-based) ultracentrifugation^110,111^.

#### *In vitro* model of hypoxia/reoxygenation

Normoxia group was washed with PBS (3x) and maintained in 0.4 mL of microtissue media at 37 °C in a humidified CO2 incubator (∼21 % O_2_). Treatment groups (IRI UT, CL2 NVs T and CERA NVs T) were washed with PBS (2x), and 1 wash with ischaemic buffer ((in mmol/L: 1.0 KH2PO4, 10.0 NaHCO3, 1.2 MgCl2.6H20, 25.0 Na(4-(2-hydroxyethyl)-1-piperazineethanesulfonic acid) (HEPES), 74.0 NaCl, 16.0 KCl, 1.2 CaCl2 and 10 mM 2-deoxyglucose, pH 6.7, gassed with pure nitrogen gas for 10 minutes). Microtissues were resuspended in 0.5 mL of ischaemic buffer and incubated in a hypoxia incubator chamber (STEMCELL Technologies, Cat # 27310) and purged with nitrogen gas for 10 minutes (99 % N_2_, 1 % O_2_). The hypoxia incubator chamber was then placed inside an incubator at 37 °C for 2 hours. Following hypoxia incubation, the ischaemic media was removed, and 0.4 mL of complete microtissue media was added with either NVs treatment (CL2 or CERA) or PBS. All groups were incubated at normal oxygen conditions (5 % CO_2_, O_2_ uncontrolled) for 24 hours.

#### Human Cardiac Troponin 1 (cTnI) ELISA

Quantification of cTnI in culture supernatant from normoxia and hypoxia/reoxygenation microtissues was performed using an ELISA (Human Cardiac Troponin 1 SimpleStep ELISA® Kit, Abcam, ab200016) according to manufacturer’s instructions. Briefly, microtissue culture media (quadruplicates per condition) was centrifuged at 2000 x g for 10 minutes to remove cell debris. 50 μL of all sample (and standards) were added to the appropriate wells in duplicate. To each well, 50 μL of the Antibody cocktail was added and incubated for 1 hour at room temperature on a plate shaker set to 400 rpm. All wells were washed three times with 1x Wash Buffer PT. Next, 100 μL of TMB Development Solution was added to each well and incubated for 10 minutes in the dark on a plate shaker set to 400 rpm. The reaction was stopped by adding 100 μL of Stop Solution to all the wells, mixed on a shaker for 1 minute. The optical density was then measured at 450 nm using a microplate spectrophotometer (Bio-Rad).

#### Cardiac microtissue contractility assay

The contractility of microtissues was recorded for normoxia and hypoxia, NV treated and NV untreated conditions, and analysed using MUSCLEMOTION macro on ImageJ, as described before^112^. Briefly, capture videos of all microtissues (n = 6) were taken using a high FPS camera (Olympus DP72, set to 15 frames per second) adapted to an inverted microscope (Olympus IX71), at 200x magnification. All videos were analysed using the MUSCLEMOTION macro at a speed window of 2 frames (15 FPS). For those microtissues which contraction propelled them out of the frame of evaluation, manual generation of stacks was required (as instructed on the original paper). Data was plotted on Graph Pad Prism V.10.4.2.

#### Proteomics: solid-phase-enhanced sample preparation

Normoxia microtissues proteome n = 4, hypoxia microtissues proteome n = 4, CL2 NV hypoxia proteome n = 4 and CERA NV hypoxia microtissue proteome n = 4 were lysed in 1 % v/v sodium dodecyl sulphate (SDS), 50 mM triethylammonium bicarbonate (TEAB), pH 8.0, incubated at 95 °C for 5 mins and quantified by microBCA (Thermo Fisher Scientific, Cat # 23235) as described^113^. Proteomic sample preparation was performed from 10 µg of protein extract as previously described{Poh, 2021 #148} using single-pot solid-phase-enhanced sample preparation (SP3){Hughes, 2019 #54}. Briefly, samples were reduced with 10 mM DTT at RT for 1 hr (350 rpm), alkylated with 20 mM iodoacetamide (IAA) (Sigma-Aldrich) for 20 min at RT (light protected), and immediately quenched with 10 mM DTT. A Sera-Mag SpeedBead carboxylate-modified magnetic particle mixture (hydrophobic and hydrophobic 1:1 mix, 65152105050250, 45152105050250, Cytiva) were added to each protein extract, washed in 50 % ethanol, and incubated for 10 min (1000 rpm) at RT. Beads were sedimented on a magnetic rack, supernatants removed and beads washed three times with 200 µL 80 % ethanol. Beads were resuspended in 100 µL 50 mM TEAB pH 8.0 and digested overnight with trypsin (1:50 trypsin: protein ratio; Promega, V5111) at 37 °C, 1000 rpm. The peptide and bead mixture was centrifuged at 20,000 g for 1 min at RT. Samples were placed on a magnetic rack and supernatant was collected and acidified to a final concentration of 1.5 % formic acid (FA), frozen at - 80 °C for 20 min, and dried by vacuum centrifugation for ∼1 h. Peptides were resuspended in 0.07 % trifluoroacetic acid (TFA), quantified by Fluorometric Peptide Assay (Thermo Fisher Scientific, Cat # 23290) as per manufacturer’s instructions, and samples normalised with 0.07 % TFA.

#### Proteomics: nano liquid chromatography–tandem mass spectrometry

Peptides were analysed on a Dionex UltiMate NCS-3500RS nanoUHPLC coupled to a Q-Exactive HF-X hybrid quadrupole-Orbitrap mass spectrometer equipped with nanospray ion source in positive, data-dependent acquisition mode ^116,117^. Peptides were separated using high resolution analytical nano liquid chromatography separated (1.9-µm particle size C18, 0.075 × 250 mm, Nikkyo Technos Co. Ltd) with a gradient of 2–28 % acetonitrile containing 0.1 % formic acid over 95 mins at 300 nl min-1 followed by 28-80 % from 95-98 mins at 300 nL min-1 at 55 °C (butterfly portfolio heater, Phoenix S&T). An MS1 scan was acquired from 350–1,650 m/z (60,000 resolution, 3 × 10^6^ automatic gain control (AGC), 128 msec injection time) followed by MS/MS data-dependent acquisition (top 25) with collision-induced dissociation and detection in the ion trap (30,000 resolution, 1×10^5^ AGC, 60 msec injection time, 28 % normalized collision energy, 1.3 m/z quadrupole isolation width). Unassigned, 1, 6-8 precursor ions charge states were rejected and peptide match disabled. Selected sequenced ions were dynamically excluded for 30 sec. Data was acquired using Xcalibur software v4.0 (ThermoFisher Scientific). The MS-based proteomics data and analysis parameters have been deposited into the ProteomeXchange Consortium via the MassIVE partner repository with the dataset identifier MSV000100182.

#### Proteomics: data processing and informatics/visualisation

Identification and quantification of peptides was performed using MaxQuant (v1.6.14.0) with its built-in search engine Andromeda^118^, as described^113,116,119^. Human-only (UniProt # 78,120 entries) sequence database (03/22) with a contaminants database was employed. N-terminal acetylation and methionine oxidations were set as variable modifications. False discovery rate (FDR) was 0.01 for protein and peptide levels. Enzyme specificity was set as C-terminal to arginine and lysine using trypsin protease, and a maximum of two missed cleavages allowed. Peptides were identified with an initial precursor mass deviation of up to 7 ppm and a fragment mass deviation of 20 ppm. Protein identification required at least one unique or razor peptide per protein group. Contaminants, and reverse identification were excluded from further data analysis. ‘Match between run algorithm’ in MaxQuant^120^ and label-free protein quantitation (maxLFQ) was performed. All proteins and peptides matching to the reversed database were filtered out.

Perseus^121^ and R studio were used to analyse the proteomic data and generate plots. G:Profiler, Reactome, STRING were used for enrichment analysis. Protein lists for samples were generated in Perseus (v1.6.14.0)^122^. For cell, NV, and NV-treated cell proteomes, proteins were identified at least once in two biological replicates per group. Protein intensities (maxLFQ) were log2 transformed. Statistical analysis and plots were generated using Perseus, R studio and GraphPad Prism. Principal component analysis, Pearson correlation matrix, and hierarchical clustering was performed in R studio using Euclidian distance and average linkage clustering, with missing values imputed at z-score 0 for heatmap generation only. R was used for data visualisation (ggplot2, ggpubr packages). In all instances significance was p < 0.05 unless otherwise indicated.

## Author Contributions

JL contributed to project coordination and design, experimental design, NVs generation, functional assays planning and execution, mass-spectrometry samples preparation/data generation/data analysis, bioinformatics analyses, preparation of figures, manuscript drafting, and provided intellectual input. JGL contributed to iPSC culture/maintenance, microtissue generation and culture, microtissue functional assays, contractility data analysis. JC contributed to mass-spectrometry samples preparation/data generation/troubleshooting/data analysis and interpretation, figure drafting, manuscript review. AMK contributed to microtissue samples preparation, cryo-sectioning, sample staining and endothelial cell differentiation of iPSC and imaging. RJP contributed to iPSC culture/maintenance, and cardiac differentiation of iPSC. AR contributed to project coordination, experimental design, provided intellectual input, and manuscript drafting and review. SYL contributed to microtissue culture/maintenance, microtissues assays and manuscript review. DG contributed to project development and coordination, experimental design, data analysis, manuscript drafting/review and preparation of figures. All authors discussed the results and commented on the manuscript.

## Supporting information

Supplementary Data (all)

Supplementary Figures (all)

## Acknowledgments

This work was supported by the National Heart Foundation of Australia (DG: Vanguard), NHMRC project grant (DG: #1139489, 1057741), Future Fund (DG: MRF1201805), Pankind (DG), Stafford Fox Medical Research Foundation (SYL), and the Victorian Government’s Operational Infrastructure Support Program. JL is supported by an Australian Government Training Program (RTP) scholarship, La Trobe University-Baker Heart and Diabetes Institute joint scholarship.

## Additional Information

Competing interests: None

## Supplemental Data

This article contains supplemental data.

- Supplementary Figure 1
- Supplementary Tables 1-14
- Supplementary Legends – Supplementary Figure and Table

## Data Availability Statement

Data generated or analysed during this study are included in this published article (and its supplementary information files) or available from Data Repositories. MS-based proteomics data is deposited to the ProteomeXchange Consortium via the MassIVE partner repository and available via MassIVE with identifier MSV000100182. Functional enrichment annotations were retrieved using g:Profiler (https://biit.cs.ut.ee/gprofiler/). Hierarchical clustering was performed in R and Perseus using Euclidian distance and average linkage clustering, with missing values imputed at z-score 0. R was also used for data visualisation.

---

**Supplementary Figure 1. Human cardiac microtissue hypoxia/reoxygenation model. (B)** Representative immunofluorescence imaging of human cardiac microtissues: Normoxia and hypoxia/reoxygenation conditions. Scale 500 nm. DAPI, VIM (Vimentin), cTnT (cardiac Troponin).

**Supplementary Table 1** - Proteome IDs of human cardiac organoids: Normoxia UT, IRI UT, CL2 NVs T and CERA NVs T - Log2 LFQ intensity

**Supplementary Table 2** - Imputed, normalised human cardiac organoids proteome

**Supplementary Table 3** - Z-scored statistically significant proteins between organoid groups.

**Supplementary Table 4** - Z-scored statistically significant proteins between Normoxia UT and IRI UT.

**Supplementary Table 5** - Z-scored statistically significant proteins between IRI UT, CL2 NVs T and CERA NVs T.

**Supplementary Table 6** - Z-scored statistically significant proteins between IRI UT and CL2 NVs T.

**Supplementary Table 7** - Z-scored statistically significant proteins between IRI UT and CERA NVs T.

**Supplementary Table 8** - Z-scored statistically significant proteins between CL2 NVs T and CERA NVs T.

**Supplementary Table 9** - Gene Ontology enrichment analyses of organoid proteome based on IRI UT

**Supplementary Table 10** - REACTOME pathway enrichment analyses of organoid proteome (IRI UT vs Treatments)

**Supplementary Table 11** - Gene Ontology enrichment analyses of organoid proteome - IRI UT and CL2 NVs T.

**Supplementary Table 12** - Gene Ontology enrichment analyses of organoid proteome - IRI UT and CERA NVs T.

**Supplementary Table 13** - REACTOME pathway enrichment analyses of organoid proteome - IRI UT vs CERA NVs T.

**Supplementary Table 14** - Z-scored statistically significant proteins common in CL2 NVs T and CERA NVs T (Scatter plot).

