## Supplementary figures and images for "An integrated cardiac microtissue proteome map extends therapeutic remodelling by nanovesicles"

### Supplementary Figures (all)

#### Slide 1
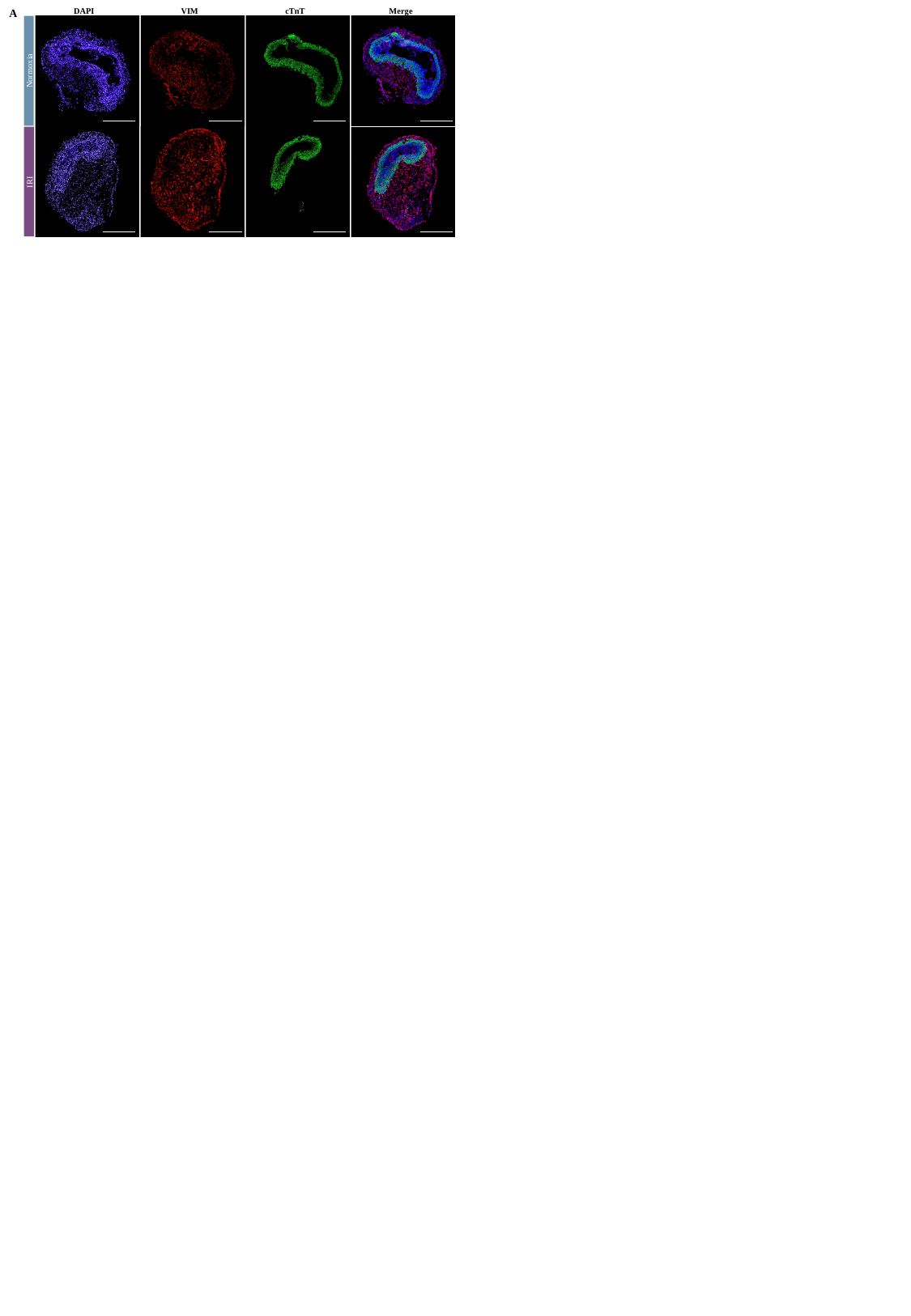

DAPI
VIM
cTnT
Merge
A
Normoxia
IRI
